# Calorie Restriction activates a gastric Notch-FOXO1 pathway to expand Ghrelin cells

**DOI:** 10.1101/2023.03.06.531352

**Authors:** Wendy M. McKimpson, Sophia Spiegel, Maria Mukhanova, Michael Kraakman, Wen Du, Takumi Kitamoto, Junjie Yu, Utpal Pajvani, Domenico Accili

## Abstract

Calorie restriction increases lifespan. While some tissue-specific protective effects of calorie restriction have been described, the impact of calorie restriction on the gastrointestinal tract remains unclear. We found increased abundance of chromogranin A+, including orexigenic ghrelin+, endocrine cells in the stomach of calorie-restricted mice. This effect coincided with increased Notch target Hes1 and Notch ligand Jag1 and was reversed when Notch signaling was blocked using the γ-secretase inhibitor DAPT. Using primary cultures and genetically-modified reporter mice, we determined that increased endocrine cell abundance was due to altered stem and progenitor proliferation. Different from the intestine, calorie restriction decreased gastric Lgr5+ stem cells, while increasing a FOXO1/Neurog3+ subpopulation of endocrine progenitors in a Notch-dependent manner. Further, calorie restriction triggered nuclear localization of FOXO1, which was sufficient to promote endocrine cell differentiation. Taken together, the data indicate that calorie restriction promotes gastric endocrine cell differentiation triggered by active Notch signaling and regulated by FOXO1.

## Introduction

Restricting calories can extend individual lifespan^1^. Initially discovered in humans, this phenomenon has been reproduced in many species, including mice^2^. Calorie restriction also improves health by delaying the onset of age-related diseases including degenerative brain disease, heart disease, type 2 diabetes, as well as certain types of cancer^3,4^. The mechanism of the alleged protective effect of calorie restriction is unknown. Interestingly, there are decreased energy expenditure and oxidative stress markers in humans whose diet was restricted by 15% over a 2 year period^5,6^. In addition, calorie restriction in humans also decreases cardiometabolic risk factors^7^ and circulating tumor necrosis factor-α^8^, a marker of inflammation.

The gastrointestinal tract is affected by calorie restriction. Enteroendocrine cells located throughout the intestine sense nutrients and releases hormones that regulate food intake and satiety^9^. Among them, ghrelin is expressed in multiple regions of the gastrointestinal tract and brain, most prominently in gastric epithelial cells^10^, and is released during food restriction to increase food intake^11^.

Differentiated intestinal cells are repopulated from stems cells, largely demarcated by Lgr5, that reside in crypts^12^. Interestingly, calorie restriction expands crypt size, while decreasing villi length^13^. In addition, it decreases the number of different cell types, including chromogranin A+ enteroendocrine cells, while increasing stem cell proliferation^14^. Fasting also promotes the fatty acid oxidation function of stem and progenitor cells^15^. The role of Lgr5+ stem cells in antral gastric tissue^16^ upon calorie restriction has not been studied. Distinct programs are activated by calorie restriction in each organ. For example, high-throughput transcriptomic, proteomic, and metabolomic analyses of liver found that caloric restriction induces RNA processing^17^. In contrast, it triggers remodeling of the skin and fur^18^. Less is known about how calorie restriction impacts cells in the stomach, although ghrelin levels reportedly increase^19^. It has also recently come to light that cells in each tissue have a unique transcription network to regulate the reversal of age-related gene expression changes by calorie restriction^20^.

Herein, we found that calorie restriction initiates a distinct gastric program affecting stem and endocrine progenitors in the stomach by modulating Notch activity. Further, during calorie restriction, FOXO1 preferentially redistributes to the nucleus in gastric endocrine progenitor cells to trigger an increase in ghrelin+ cells, thereby stimulating food intake.

## Results

### Calorie restriction alters gastric endocrine cell composition

Previous research shows that plasma ghrelin increases after calorie restriction^19^. To investigate whether this is due to increased ghrelin cell number, we calorie restricted FOXO1-Venus (FoxV) knock-in mice^21^ by 30% of their daily food intake (CR), whereas controls had unlimited access to food (ad libitum). After calorie restriction, FoxV CR mice showed decreased weight (Figure 1A), fasting glucose (Figure 1B), and lean mass (Figure 1C) compared to controls. The latter gained weight and lean mass, as expected (Figures 1A and 1C).

**Figure 1:**
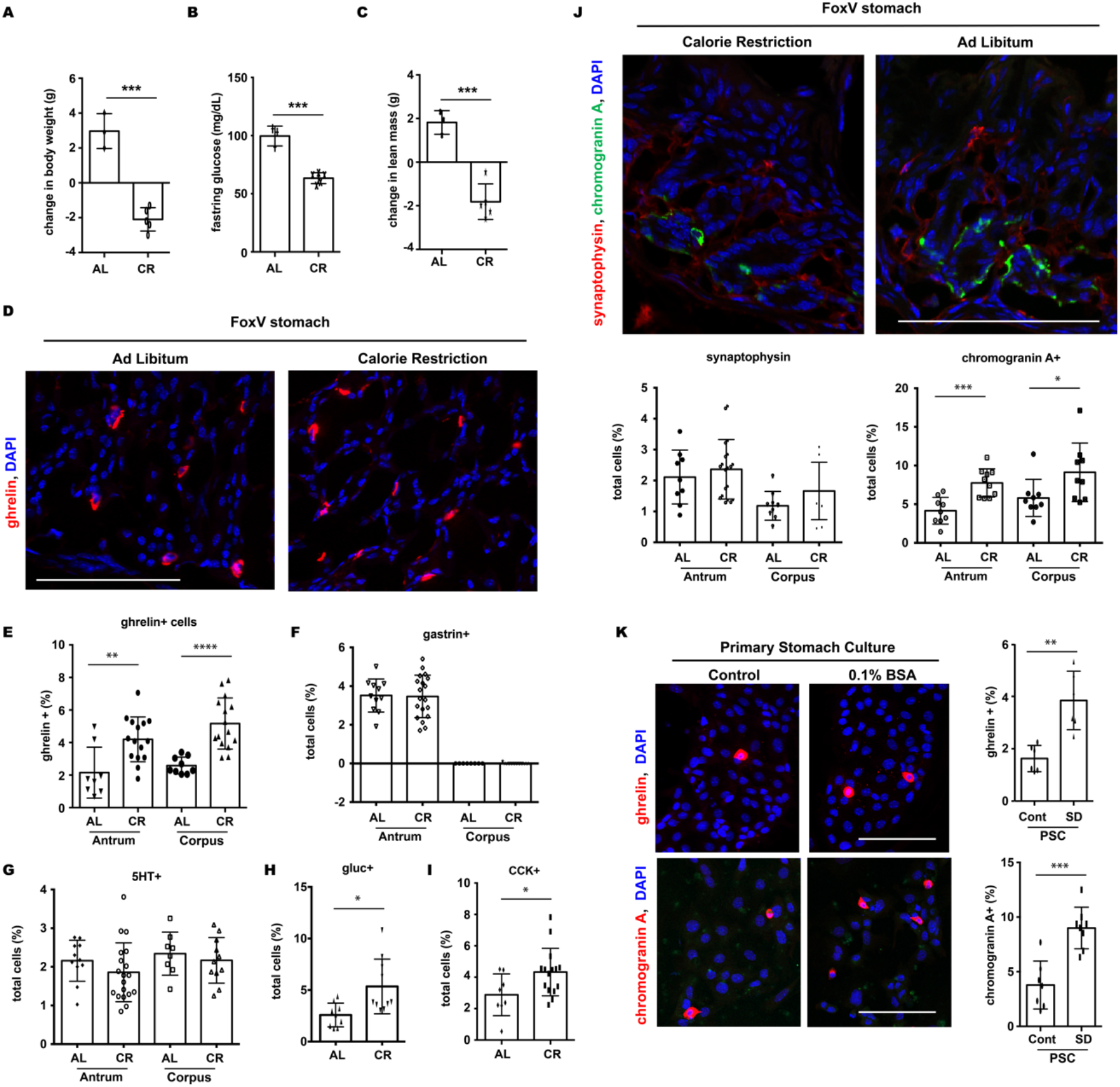
Gastric endocrine cells become more abundant after calorie restriction. A) Change in body weight after FOXO1-Venus reporter mice (FoxV) were fed ad libitum (AL, n = 3 mice) or 30% food restricted (CR, n=5 mice) for 4 weeks. B) Blood glucose measurements after a 5 h fast in AL and CR mice. C) Change in lean mass measurements in AL and CR mice. D) Fluorescent staining of ghrelin-expressing cells from stomach tissue of AL and CR FoxV mice. E) Quantification of panel D. F) - I) Quantification of endocrine cell populations in AL and CR FoxV stomach. J) Immunofluorescence (top) with quantification (bottom) for Synaptophysin and Chromogranin A endocrine markers in FoxV mice after CR. K) Immunohistochemistry (left) with quantification (right) of primary stomach cultures (PSC) generated from wild type mice and cultured in control and serum-deprived (0.1% BSA) media and stained for endocrine markers. Scale bar is 100μm. DAPI counterstains nuclei. Bar graphs show mean ± SD. Each dot represents values from individual mouse or quantification of a microscopic field from PSC or AL and CR FoxV mice. * P < 0.05, ** P < 0.01, *** P < 0.001, and **** P < 0.0001.

We assessed ghrelin+ cell number by immunostaining and found a doubling of cell abundance in both antrum and corpus (Figure 1D and 1E). To determine if other endocrine cell populations were altered, we immunostained stomach sections for 5HT and gastrin and found no changes (Figures 1F and 1G). We saw a slight, but significant increase in cells positive for glucagon and CCK (Figures 1H and 1I). We next examined alterations in differentiated endocrine cells after calorie restriction using general endocrine cell markers. While the number of synaptophysin+ cells did not change, we found a significant increase of chromogranin A+ cells in antrum and corpus (Figures 1J). To recapitulate these changes *ex vivo*, we cultured primary stomach cultures (PSC) in serum-free medium and found a similar induction of ghrelin+ and chromogranin A+ cells (Figure 1K).

### Notch signaling is triggered by food restriction

To determine the signaling mechanisms regulating changes in endocrine cell abundance with calorie restriction, we next surveyed molecular pathways involved in gastric cell differentiation. We found a substantial increase in cells positive for the Notch transcriptional target Hes1 after calorie restriction, consistent with increased Notch activity (Figure 2A). Importantly, We also saw increased Notch receptor levels in antral cells after calorie restriction (Figure 2B). Further, culturing primary stomach cells in medium supplemented with bovine serum albumin mimics the increase in Notch activity seen during calorie restriction (Figure 2C).

**Figure 2:**
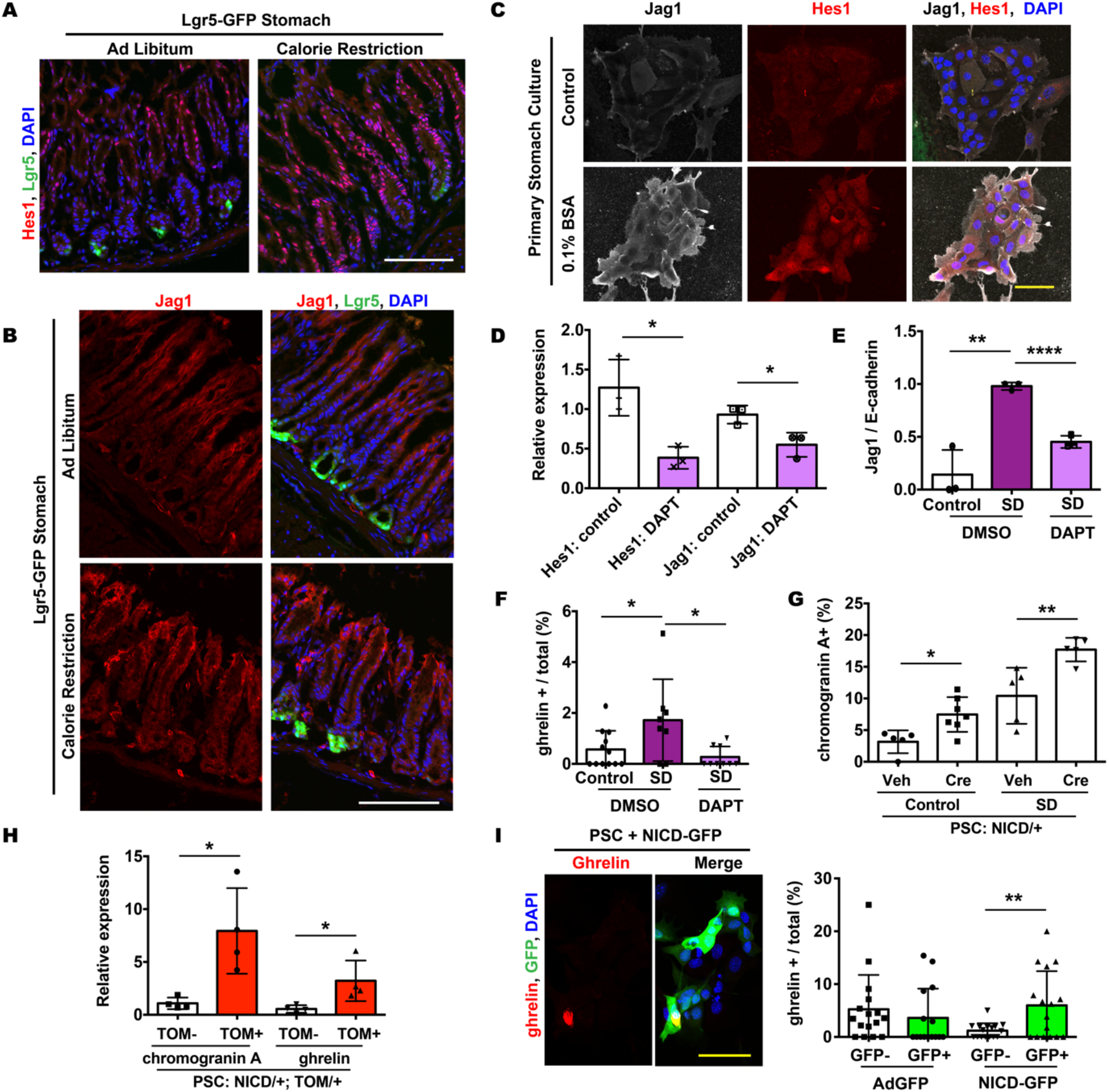
Calorie restriction activates Notch signaling to increase Ghrelin+ cells. A) and B) Immunohistochemistry of antral stomach tissue from Lgr5-GFP reporter after calorie restriction. C) Immunohistochemistry of primary stomach cultures (PSC) generated from wild type mice and cultured in control or serum-deprived (SD; 0.1% BSA) media. D) Gene expression of PSC after overnight treatment with γ-secretase inhibitor DAPT (10μM). E) Gene expression of PSC grown in control, serum-deprived (0.1% BSA) media, or serum-deprived media treated with DAPT. F) Quantification of ghrelin immunohistochemistry of wild type PSC grown in control, serum-deprived (0.1% BSA) media, or serum-deprived media treated with DAPT. G) Quantification of chromogranin A staining in PSC generated from NICD/+ mice and treated with vehicle (Veh) or 4μM recombinant TAT-cre (Cre) for 1 day. PSC were stained following an overnight recovery after Cre treatment. H) Gene expression of cells purified by FACS. PSC analyzed were from NICD/+; Tomato(TOM)/+ mice and treated overnight with TAT-cre to induce NICD. Cells collected were either negative (TOM-) or positive (TOM+) for endogenous tomato, which fluoresces upon cre recombination. I) Representative image (left) with quantification (right) of wild type primary stomach cultures transduced overnight with control (AdGFP) or NICD (NICD-GFP) adenovirus and then immunostained for Ghrelin. White scale bar is 100μm, yellow scale bar is 50μm. DAPI counterstains nuclei. Gene expression is normalized to E-cadherin. Bar graphs show mean + SD. Each dot represents quantification of a microscopic field. * P < 0.05, ** P < 0.01, and **** P < 0.0001.

To determine whether the increase in Notch-active cells during calorie restriction is linked to the increased ghrelin cells, we incubated PSC with the γ-secretase inhibitor DAPT, which blocks Notch signaling (Figure 2D and 2E). Interestingly, the increased number of ghrelin+ cells in PSC caused by serum deprivation was reversed by overnight treatment with DAPT (Figure 2F). Conversely, cre recombination-mediated expression of the Notch intracellular domain (NICD) triggered an increase of chromogranin A-immunoreactive cells (Figure 2G). Cells overexpressing NICD also contained increased chromogranin A and ghrelin gene expression (Figure 2H). Adenoviral-mediated expression of NICD also increased ghrelin+ cells (Figure 2I). These data suggest that activation of Notch signaling is necessary and sufficient to increase chromogranin A+ and ghrelin+ cells during calorie restriction.

### Active Notch perturbs gastric stem cells

Notch signaling regulates gastric Lgr5 stem cell differentiation^22^. To determine if changes in differentiated endocrine cells were due to changes in stem and progenitor populations, we calorie restricted Lgr5-GFP and Neurog3-GFP knock-in reporter mice to label stem and endocrine progenitor cells, respectively. These reporter mice had lower glucose levels (Figure 3A) and decreased lean mass (Figure 3B) upon calorie restriction. In the stomach, calorie restriction triggered a decrease of Lgr5-GFP+ cells (Figures 3C and 3D). In contrast, cells that express transcription factor Sox2, a marker of multipotent stomach stem cells^23^, were increased (Figure 3E).

**Figure 3:**
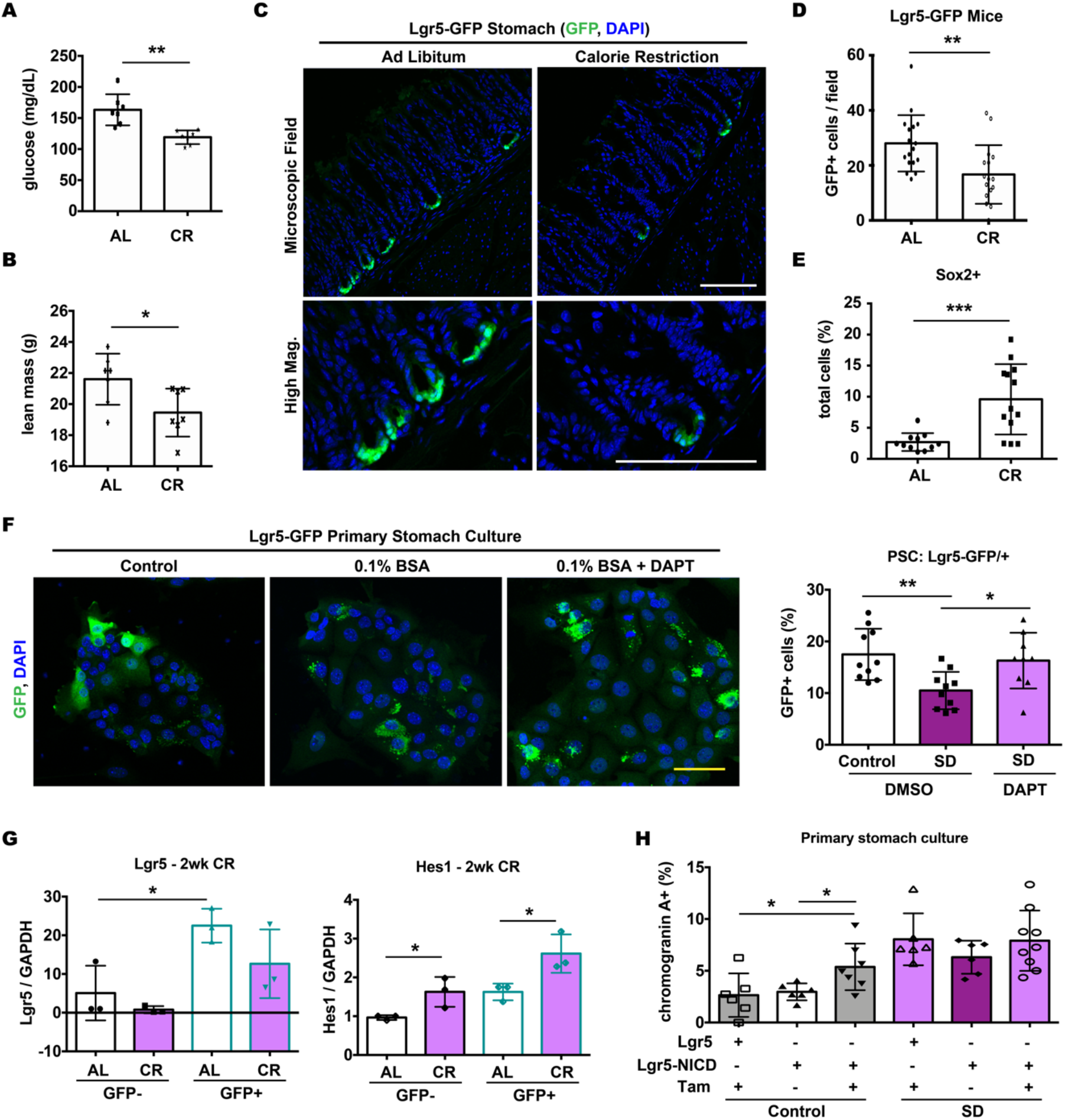
Calorie restriction decreases the number of gastric Lgr5+ stem cells in a notch-dependent manner. A) Fed blood glucose measurements after Lgr5-GFP and Neurog3-GFP reporter mice were fed ad libitum (AL) or 30% food restricted (CR) for 4 weeks. B) Lean mass measurements in AL and CR mice. C) Stomach tissue from Lgr5-GFP reporter mouse showing endogenous fluorescence in AL and CR tissue. D) Quantification of GFP+ cells in Lgr5-GFP mice shown in panel C. E) Quantification of Sox2+ cells in stomach tissue from FOXO1-Venus reporter mouse (FoxV) after CR. F) Immunohistochemistry (left) with quantification (right) of primary stomach cultures (PSC) generated from Lgr5-GFP mice stained for GFP. PSC were grown in control or serum-deprived (SD; 0.1% BSA) media and treated overnight with vehicle (DMSO) or γ-secretase inhibitor DAPT (10μM). G) Gene expression of cells purified by FACS. PSC analyzed were from Lgr5-GFP mice that were fed ad libitum (AL) or 40% food restricted (CR) for 2 weeks. Cells collected were either negative (GFP-) or positive (GFP+) for endogenous Lgr5-GFP. H) Quantification of chromogranin A immunohistochemistry in PSC generated from mice heterozygous for Lgr5-GFP reporter allele (Lgr5) or from mice heterozygous for both Lgr5-GFP and NICD. PSC were cultured in control or SD media and treated overnight with 4-Hydroxytamoxifen (Tam, 1μM) to activate cre recombinase from Lgr5 allele. White scale bar is 100μm, yellow scale bar is 50μm. DAPI counterstains nuclei. Bar graphs show mean + SD. Each dot represents an individual mouse measurement (panels A and B) or quantification of a microscopic field from PSC or AL (Lgr5-GFP mice: n=3) and CR (Lgr5-GFP mice: n=3) mice. * P < 0.05, ** P < 0.01, *** P < 0.001.

To test the involvement of Notch signaling in stem cell maintenance after calorie restriction, we used PSC. There were 40% fewer Lgr5-GFP+ cells when PSC isolated from Lgr5-GFP mice were cultured in serum-deprived compared to serum-containing medium (Figure 3F).

This decrease was dependent upon Notch activation, as treatment of PSC with DAPT restored the number of Lgr5-GFP+ cells (Figure 3F). Interestingly, both Lgr5-GFP– and Lgr5-GFP+ cells had increased levels of the Notch transcription factor Hes1 after calorie restriction (Figure 3G). We leveraging the tamoxifen-inducible cre cloned in the Lgr5 locus of Lgr5-GFP mice to test the effect of Notch activation in Lgr5+ cells on endocrine cell abundance. For this, we isolated PSC from Lgr5-GFP/+; NICD/+ mice and treated them with tamoxifen to induce NICD expression. Activation of NICD in Lgr5-expressing cells increased the number of chromogranin A+ cells (Figure 3H). Importantly, this phenotype was seen in PSC grown in control medium and numbers chromogranin A+ cells were similar to those cultured in serum-deprived conditions.

### Notch promotes expansion of FOXO1+, Neurog3+ endocrine progenitors

Using the calorie restricted reporter mice described above (Figure 3A and 3B), we next analyzed endocrine progenitor cell populations. Neurog3-GFP+ endocrine progenitor cells increased in the stomach after calorie restriction (Figures 4A and 4B). Similar to *in vivo*, PSC grown in serum-deprived medium had more Neurog3-GFP+ cells as detected by endogenous fluorescence using FACS (Figure 4C and 4D). Similar to previous reports^24,25^, Neurog3-GFP+ cell were enriched for the transcription factor FOXO1 (Figure 4E). Serum-deprivation also increased Neurog3-GFP+ cells by 180% as detected by immunohistochemistry (Figure 4F). This increased abundance was reversed when cultures were treated with the Notch inhibitor DAPT (Figure 4F).

**Figure 4:**
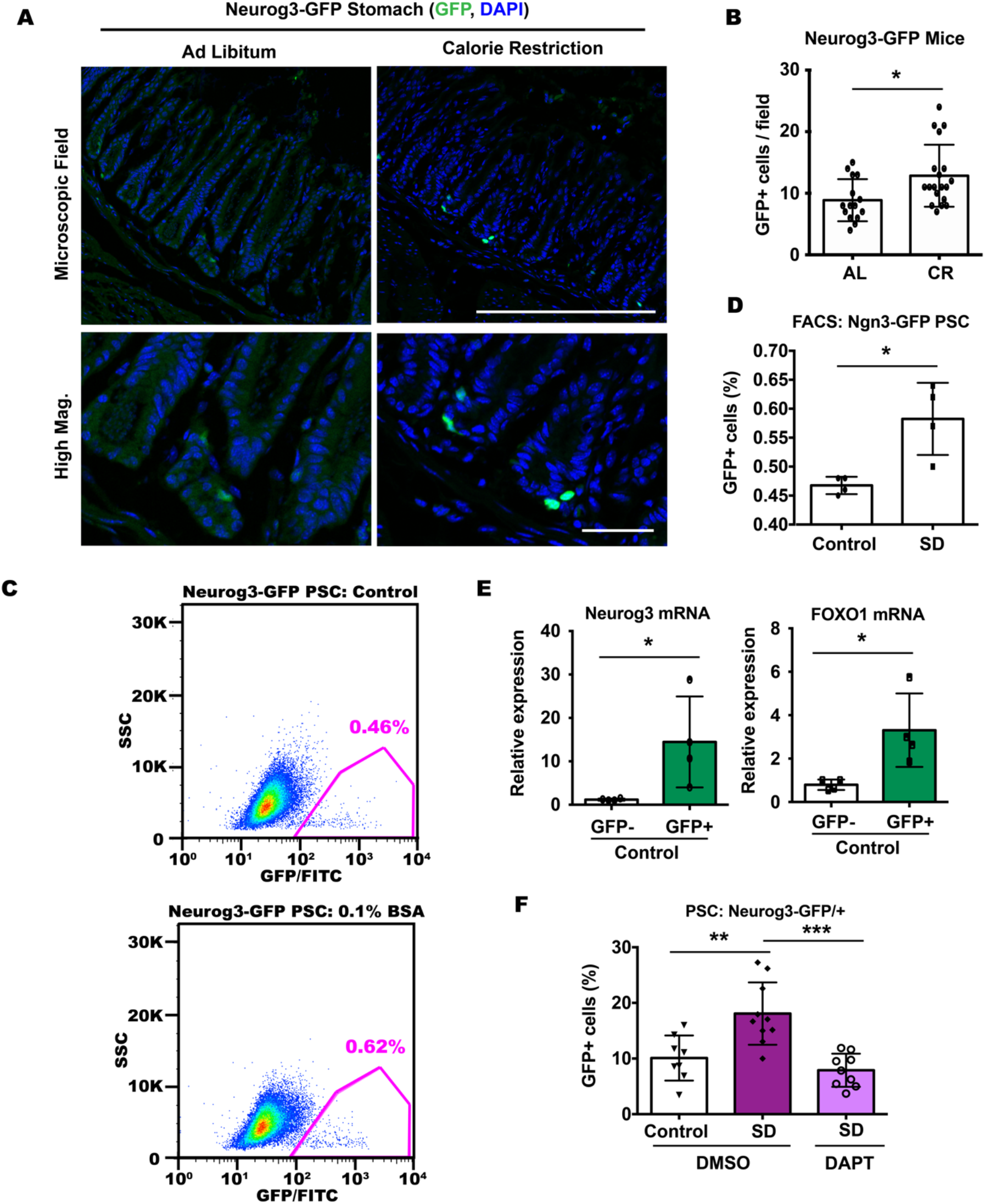
The abundance of endocrine progenitor cells is increased by calorie restriction through altered notch signaling. A) Stomach tissue from Neurog3-GFP/+ reporter mice fed ad libitum (AL) or 30% food restricted (CR) for 4 weeks. Endogenous GFP fluorescence is shown. Metabolic characterization is displayed in Figure 3A and 3B. B) Quantification of GFP+ cells in Neurog3-GFP+ mice shown in panel A. C) Representative FACS dot plots of primary stomach cells isolated from Neurog3-GFP/+ mice after overnight culture in control or serum-deprived (SD; 0.1% BSA) media. GFP+ population is outlined in pink. D) Quantification of panel C. E) Gene expression of GFP+ cells purified by FACS. PSC analyzed were from Neurog3-GFP mice after overnight culture in control media shown in panel C. F) Quantification of primary stomach cultures (PSC) generated from Neurog3-GFP mice immunostained for GFP cultured in control or SD media and treated overnight with vehicle (DMSO) or γ-secretase inhibitor DAPT (10μM). White scale bar is 100μm. DAPI counterstains nuclei. Bar graphs show mean + SD. Each dot represents quantification of a microscopic field from PSC for AL (Neurog3-GFP mice: n=4) and CR (Neurog3-GFP mice: n=4) mice. * P < 0.05, ** P < 0.01, *** P < 0.001.

Since FOXO1 is expressed in Neurog3+ cells^24,25^, we next analyzed the effect of calorie restriction on FOXO1-expressing cells in the stomach. To detect FOXO1, we used GFP immunohistochemistry in FoxV mice^25^, and used pancreatic islets as a positive control (Figure 5A). No reactivity was detected in wild type mice (Figure 5B). Immunohistochemistry revealed that stomachs of calorie-restricted mice had ~135-170% more FoxV+ cells in a region-specific manner in the antrum and corpus (Figure 5C – 5E). FOXO1-expressing Neurog3+ cells also increased after calorie restriction (Figure 5F).

**Figure 5:**
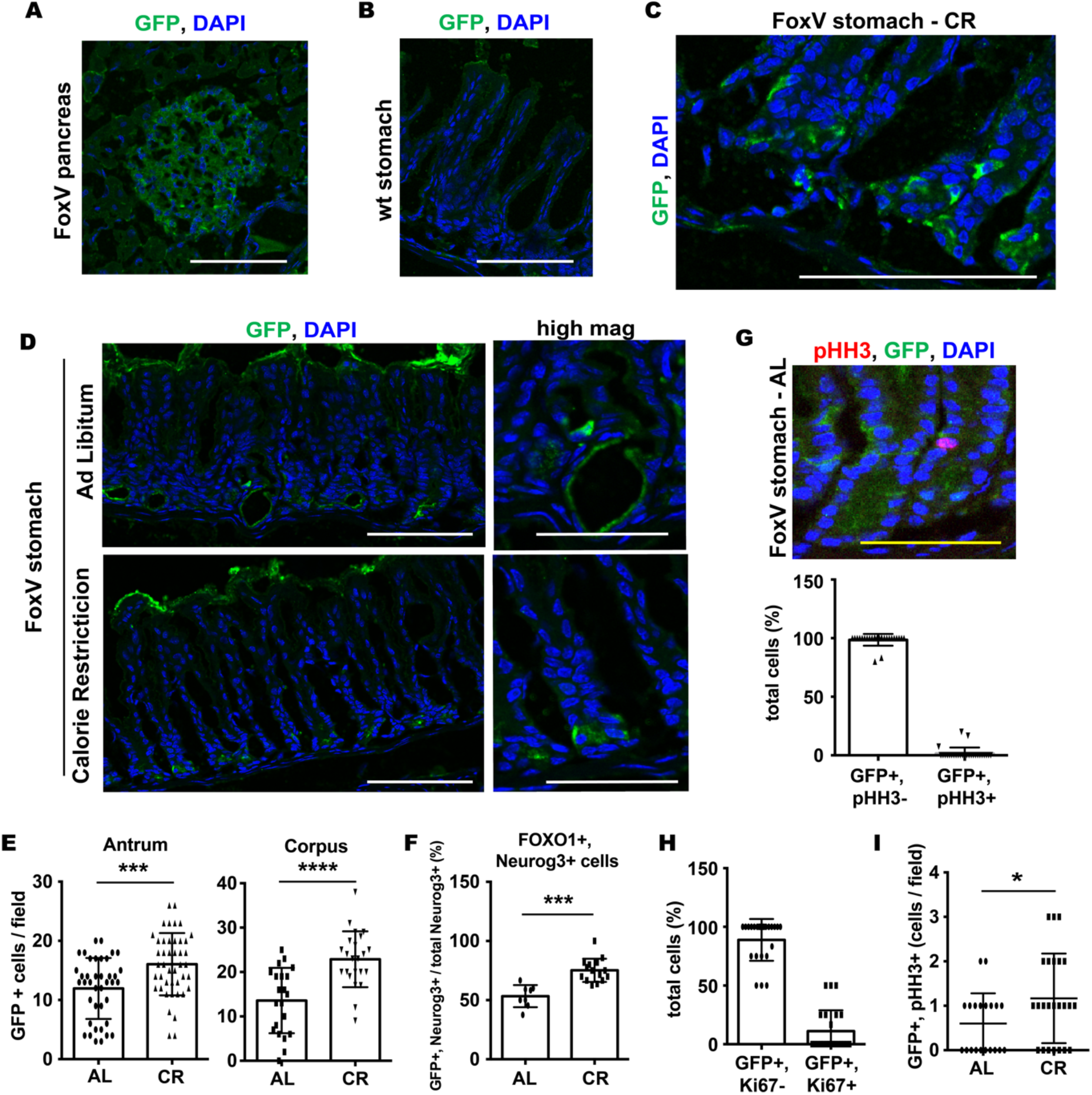
There are more gastric endocrine progenitor FOXO1+ cells after calorie restriction. A) Pancreatic tissue from FOXO1-Venus reporter mouse (FoxV) stained for GFP (which recognizes Venus reporter). This represents a positive control. B) Wild type stomach tissue stained from GFP showing no positive signal in stomach tissue lacking FoxV reporter. C) and D) Representative images of GFP staining (which recognizes FOXO1-Venus reporter) in FoxV stomach of mice after 4 week calorie restriction (CR, n=5 mice) or in control mice fed ad libitum (AL, n = 3 mice). E) Quantification of GFP+ cells in AL and CR mice in the antrum (left) and corpus (right) stomach. F) Quantification of GFP (FoxV+ cell) and Neurogenin3 (Neurog3) colocalization in FoxV AL and CR mice. G) Immunostaining (top) with quantification (bottom) of GFP colocalization with proliferation marker phosphorylated histone H3 (pHH3) in the stomach of AL FoxV mice. H) Quantification of GFP colocalization with a second proliferation marker Ki67 in the stomach of AL FoxV mice. I) In a separate experiment, quantification of GFP colocalization with the pHH3 in stomach of AL and CR FoxV mice. White scale bar is 100μm, yellow scale bar is 50μm. DAPI counterstains nuclei. Bar graphs show mean + SD. Each dot represents a quantification of a microscopic field from AL and CR mice. * P < 0.05, *** P < 0.001, **** P < 0.0001.

Changes in cell number can be due to cell proliferation. Interestingly, under basal conditions, FOXO1 + stomach cells are minimally proliferative, as indicated by the paucity of GFP+ cells colocalizing with proliferation markers phospho-histone H3 (pHH3) (Figure 5G) and Ki67 (Figure 5H). However, there was a significant 195% increase in Foxo1-expressing GFP+, pHH3+ cells after calorie restriction (Figure 5I).

### Nuclear FOXO1 triggers increase in Ghrelin+ cells

In ad libitum-fed conditions, FOXO1 is mainly detected in the cytoplasm of cells and rarely overlaps with the nuclear marker DAPI (Figure 6A). Interestingly, there were regions in the stomach of calorie restricted mice where GFP staining also noticeably overlapped with the nuclear marker DAPI (Figure 6B). Importantly, such regions were not found in mice fed ad libitum. This staining pattern is also in contrast with other regions where GFP staining was either exclusively cytoplasmic or diffused to the entire cell (Figure 5C).

**Figure 6:**
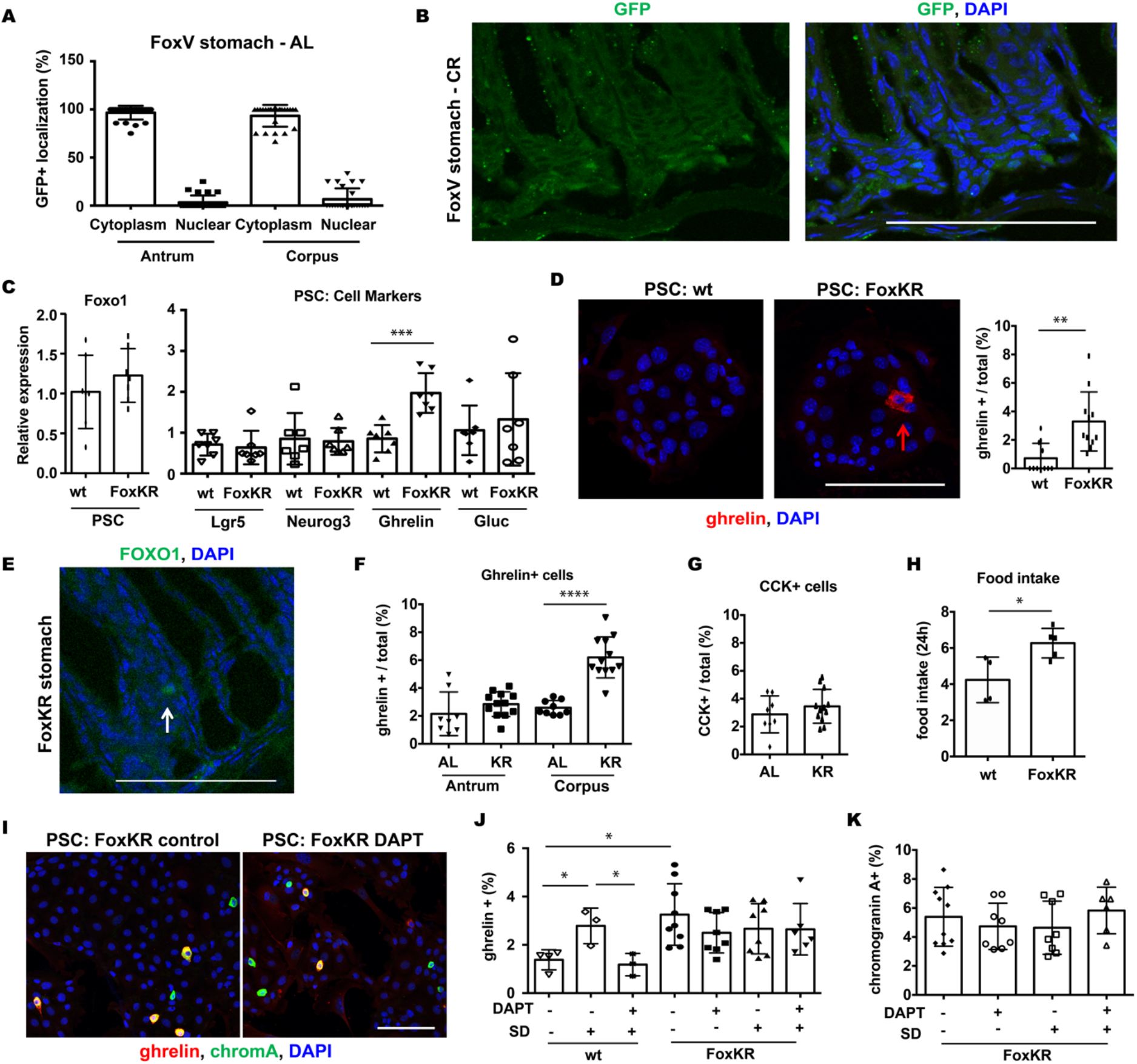
Nuclear FOXO1 promotes differentiation of Ghrelin cells in primary stomach culture and in mice. A) Quantification of subcellular localization of GFP staining in FOXO1-Venus reporter (FoxV) mice stained for GFP (which recognizes Venus reporter) that have been fed ad libitum (AL) FoxV mice. Nuclear represents colocalization of GFP with DAPI. B) Representative image showing region with nuclear GFP staining in FoxV stomach after calorie restriction (CR). C) Gene expression in primary stomach cultures (PSC). Primary stomach cells were isolated from wild type (wt) mice and mice homozygous for a constitutively active FOXO1 mutant (FoxKR) that localizes to the nucleus. Stomach markers include: neurogenin3 (Neurog3) and glucagon (Gluc). D) Representative image (left) with quantification (right) of PSC from indicated mice stained for Ghrelin. E) Florescent staining for FOXO1 in FoxKR stomach. F) and G) Quantification of ghrelin+ (panel F) and CCK+ (panel G) cells in control (AL) and FoxKR (KR) stomach. AL (n=3 mice) bars represent the same values quantified in figure 1 and KR (n=4 mice). H) Daily food intake measurements for wild type (wt) (n=4) and KR (n=5) mice. Values presented represent an average 24h intake measured over two consecutive days. I) Immunohistofluorescence of primary stomach cultures (PSC) generated from KR mice cultured in control media after overnight treatment with vehicle (control) or γ-secretase inhibitor DAPT (10μM). J) Quantification of ghrelin+ cells in PSC from wt and KR mice grown in control or serum-deprived (SD; 0.1% BSA) media and treated overnight with DAPT. K) Quantification of chromogranin A+ cells in FoxKR PSC at the indicated conditions. Scale bar is 100μm. DAPI counterstains nuclei. Bar graphs show mean + SD. Each dot represents quantification of a microscopic field. * P < 0.05, ** P < 0.01, *** P < 0.001, and **** P < 0.0001.

FOXO1 actively cycles between nucleus and cytoplasm. To test the role of nuclear FOXO1 in gastric endocrine cell remodeling following calorie restriction, we studied PSC from knock-in mice homozygous for a constitutively nuclear acetylation defective FOXO1 (FoxKR)^25^. As expected, *Foxo1* mRNA levels were similar in PSC created from wild type and FoxKR mice (Figure 6C). While expression of different stomach cell markers, including *Lgr5* and *Neurog3*, was minimally altered under ad libitum-fed conditions, *ghrelin* mRNA levels were significantly increased (Figure 6C). Ghrelin+ cells were also more abundant in PSC from stomach cells isolated from FoxKR mice compared to controls (Figure 6D).

We confirmed these results *in vivo* using immunohistochemistry. Importantly, FOXO1 was indeed nuclear in FoxKR mice (Figure 6E). While the number of ghrelin+ cells in the antrum of these mice was minimally changed from controls, there was a striking increase in ghrelin+ cells in the corpus (Figure 6F). The abundance of CCK+ cells was not altered in FoxKR mice (Figure 6G). FoxKR mice also had increased food intake (Figure 6H). Taken together, our results suggest a role for nuclear FOXO1 in promoting the differentiation of ghrelin+ cells with calorie restriction.

To test whether the role of FOXO1 was dependent on Notch signaling, we treated PSC from wild type and FoxKR mice with DAPT and measured chromogranin A+ and ghrelin+ cells by immunohistochemistry (Figure 6I). PSC generated from FoxKR stomachs had an increased number of ghrelin+ cells compared to wild type PSC (Figure 6J). And while there were more ghrelin+ cells in PSC from wild type mice upon serum-deprivation, and this was reversed by DAPT treatment, neither treatment affected the number of ghrelin+ cells in FoxKR PSC (Figure 6J). Similarly, serum-deprivation and DAPT treatment did not alter the abundance of chromogranin A+ cells in FoxKR PSC (Figure 6K). These data link nuclear FOXO1 with ghrelin cell differentiation downstream of Notch activation.

## Discussion

We found that, different from what is seen in intestine, calorie restriction triggers increases in differentiated endocrine and Neurog3+ progenitor cells through increased Notch signaling in gastric stem and progenitor cells. We further show, using an active FOXO1 mutant, that FOXO1 localizes to the nucleus of Neurog3+ cells with calorie restriction, and that this is associated with increased numbers of ghrelin+ cells.

Our data demonstrate that ghrelin cell numbers increase upon calorie restriction. While we do not know whether ghrelin is actively produced and secreted, this is likely in view of the increase in plasma ghrelin during fasting^26,27^. A link between ghrelin and calorie restriction has been suggested based on studies showing increased ghrelin levels with calorie restriction^19^. Interestingly, ghrelin may also have a protective effect against aging^28,29^. It is not clear whether the increase in ghrelin+ cells is an adaptive change due to decreased food consumption, or a protective effect of calorie restriction itself.

Mechanistically, our data suggest that the increase in ghrelin+ cell number is dependent upon active FOXO1 signaling. FOXO1 regulates cellular differentiation and growth^30^, and has been found in Neurog3+ progenitor and 5HT+ cells in the large and small intestine of mice and humans^24,31^. We have recently characterized a relatively rare population of FOXO1+ cells in the stomach^25^, colocalizing with markers of acid-secreting parietal as well as Neurog3+ cells. In these cells, FOXO1 regulates transcription of the gene encoding cyclin E1 (CCNE1). Our data suggest that, under calorie restriction, FOXO1 marks an expanding subpopulation of Neurog3+ cells. Interestingly, FOXO1 is found in the nucleus of these cells and this is associated with increased numbers of ghrelin differentiation program. Evidence suggests that FOXO1 mediates changes occurring with calorie restriction in other tissues as well. For example, FOXO1 haploinsufficiency is linked to reduced incidence of tumors with aging, although there was no effect on lifespan^32^.

FOXO1 also has been shown to have a role in maintaining telomere size during calorie restriction in the heart^33^.

Here we demonstrated that calorie restriction alters the abundance of stomach epithelial stem and progenitor cells by increasing gastric Notch signaling, specifically Jag1 and Hes1. It remains to be determined what triggers Notch activation. Stem and progenitor cells in different tissues are sensitive to diet. For example, fasting promotes the function of stem and progenitor cells in the intestine by inducing fatty acid oxidation^15^. Prolonged fasting also promotes selfrenewal of hematopoietic stem cells^34^. Conversely, mice fed a high-fat diet have increased numbers of intestinal stem cells through a peroxisome proliferator-activated receptor delta-dependent program^35^. Interestingly, a low-lipid diet activates Notch signaling to influence cell differentiation in the Drosophila intestine^36^. Given the metabolic roles of FOXO1, it will be interesting to determine expression profiles of FOXO1-expressing Neurog3+ cells in response to diet.

Notch regulates stomach Lgr5+ cell proliferation^22^ through a DLL1-expressing niche cell located in the antral gland base to blunt their differentiation^37^. Notch inhibition generally promotes differentiation. It is unclear why the opposite result is seen in the stomach after calorie restriction, but it may indicate an effect on specific stem and progenitor cell subpopulations. In support of this, while Lgr5+ stem cells were decreased with calorie restriction, we saw a striking increase in Sox2+ cells, a distinct multipotent stem cell population in the stomach^23^. A different program may also be activated in gastric stem cells under calorie restriction. Notch-regulated Lgr5 stem cell maintenance is associated with increased mTOR signaling^22^. While we did not explore alterations of mTOR, calorie restriction modulates the function of intestinal stem cells through mTORC1 in Paneth cells, a component of the intestinal stem cell niche^13^. This expansion is dependent on mTORC1 and SIRT1^14^. In mice, calorie restriction also preserves the function of a pool of reserve stem cells in the intestine by inhibiting mTORC1^38^.

## Methods

### Mice

FoxV mice were generated as previously described^21^. Neurog3-GFP^39^ and FoxKR^40^ mice were previously described, and Lgr5-GFP, NICD, and Tomato mice were purchased from Jackson laboratories. Unless it is otherwise indicated, tissue was collected from mice 10-12 weeks of age. Male and female mice were used in equal numbers. All studies were approved by the Columbia University Institutional Animal Care and Utilization Committee.

### Calorie Restriction

Mice were fed a standard chow diet (SCD) (13.1%, 62.4% and 24.5% calories from fat, carbohydrate and protein respectively; PicoLab Rodent Diet 20, 5053; Purina Mills). All mice were acclimated to being singly housed for a week with individual food intake determined over the last 3 days. Thereafter mice were assigned to groups with an equal starting body mass and either had access to food and water *ad libitum* (Ad Lib), were calorie-restricted by 30% with food placed into the cage twice daily for 4 weeks, or were calorie-restricted by 40% with food placed into the cage twice daily for 2 weeks as indicated in the figure legends. CR mice received 2.6 – 2.8 g/day of SCD per mouse. Blood glucose was measured after 5h fast using a OneTouch glucose monitoring system (LifeScan). Lean mass was measured by functional MRI.

### Fluorescent Imaging

After dissection, stomach tissue was fixed in 4% paraformaldehyde (PFA, Electron Microscopy Sciences, cat no. 15710) for 1.5h on ice. Tissue was subsequently place in 30% sucrose (in 1xPBS) overnight, embedded in OCT, flash frozen, and sectioned at 5 μM for staining. For primary cultures, cells were grown on gelatin-coated chamber slides (Corning, cat. no. 354114), fixed in 4% paraformaldehyde for 10m at room temperature and permeabilized in PBS with 0.1% Triton X-100 (Acros Organics, Cat. no. 42235-5000) for 10min.

For immunohistochemistry, tissue was blocked in 10% goat (Vector, Cat. no. S-1000) or donkey (Jackson Immuno Research Labs, cat. no. 017000121) serum for 1 hour at room temperature and then incubated with primary antibodies diluted in blocking solution overnight. The primary antibodies used were: GFP (Molecular Probes, cat no. A-6455, 1:200), GFP (Abcam, cat no. ab6662, 1:100), Ghrelin (Novus Biologicals, cat no. MAB8200, 1:100), Ki67 (Abcam, cat no. ab16667, 1:50), Neurogenin3 (Abcam, 1:50, cat. no. ab38548), Sox2 (Abcam, cat no. ab97959, 1:200), Chromogranin A (Abcam, cat no. ab45179, 1:200), 5HT (Novus Biologicals, cat. no. NB120-16007, 1:50), Gastrin (Abcam, ab232775, 1:200), Glucagon (Takara, cat no. M182, 1:500), CCK (Thermo Fisher, PA5-32348, 1:400), Synaptophysin (Millipore, cat no. MAB5258, 1:200), Jag1 (R and D, cat no. AF599, 1:100), Hes1 (Santa Cruz, cat no. sc25392, 1:100), and pHH3 (Abcam, cat no. ab5176, 1:100). Alexa Fluor 488, 568, and 647 (Molecular Probes, dil. 1:1000) secondary antibodies of corresponding species were used to detect primary antibodies. All secondaries were diluted 1:1000 in blocking serum and incubated for 1h at room temperature. Cover slips were applied using VECTASHIELD Vibrance Antifade Mounting Medium with DAPI (Vector Laboratories, cat no. H-1800-10) to counterstain nuclei. Images were obtained using a Zeiss LSM 510/710 confocal microscope (Zeiss).

### Primary stomach culture

Primary stomach cells were isolated from the indicated mice as previously described^25^. Briefly, after serosa was removed, the antral stomach was incubated with digestion buffer (5mM EDTA (Invitrogen, cat. no. 15575-038) in 1xPBS) at 4°C for 1.5h. Following a second digestion at 37°C for 30m in TrypLE (Invitrogen, cat. no. 12604013) supplemented with 2 U/μl DNAse-1 (Sigma-Aldrich, cat. no. 10104159001) and 10 μM Y-27632 (Rock Inhibitor, Sigma-Aldrich, cat. no. Y0503-1MG), cells were washed in ice-cold basic media (Advanced DMEM/F12 (Invitrogen, cat. no. 12634010) supplemented with 10mM HEPES (Invitrogen, cat. no. 15630080), 1 x Glutamax (Invitrogen, cat. no. 35050061), 1% pen/strep (Gibco, cat. no. 15140-122), and 1% bovine serum albumin, fraction V (Fisher, cat. no. BP1600-100)) and passed through a 40μM cell strainer (Fisher, cat. no. 22363547).

For 2-D culture, primary stomach cells were plated on mouse embryonic fibroblasts (Gibco, cat. no. A34180) at density .5×10^6^ cells / 6-well plate coated with gelatin (Millipore, cat. no. ES-006-B). Cells were cultured in basic culture media (described above) supplemented with 10 μM rock inhibitor, 1 x N2 supplement (Invitrogen, cat. no. 17502048), 1 x B-27 supplement (Invitrogen, cat. no. 17504044), 1 mM N-Acetyl-L-cysteine (Sigma-Aldrich, cat. no. A9165-5G), and 1 x Primocin (Invitrogen, cat. no. NC9141851). Rock inhibitor was removed from media after overnight recovery. Media for serum-deprivation conditions was modified to contain 0.1% bovine serum albumin, fraction V. Unless otherwise indicated, cells were cultured overnight at serum-deprived conditions. Where indicated, cells were treated with 10 μM DAPT (N-[N-(3, 5-difluorophenacetyl)-l-alanyl]-s-phenylglycinet-butyl ester) and 4 μM TAT-cre (Millipore, cat. no. SCR508) overnight.

### FACS

After culture, primary stomach cells were collected using trypsin and resuspended in basic culture media. Cells were sorted using a BD Influx with a blue (488-nm) laser and 530/40 band pass filter to detect endogenous GFP fluorescence and a green (561-nm) laser and 610/20 band pass filter to detect tomato fluorescence and analyzed using FlowJo software (BD Life Sciences).

### qPCR

RNA was isolated from primary stomach cultures using an Arcturus PicoPure RNA Isolation Kit (Applied Biosystems, cat no. KIT0204) as recommended by the manufacturer with the addition of the optional 15 minute DNaseI treatment. cDNA was generated from RNA using qScript cDNA SuperMix (Quanta Biosciences, cat no. 101414-106) as previously described^41^. qPCR reactions were completed using GoTaq qPCR Master Mix (Promega) with the following primer pairs: *Hes1:* forward: CACTGATTTTGGATGCACTTAAGAAG, reverse: CCGGGGTAGGTCATGGCGTTGATCT; *Jag1:* forward: CCTCGGGTCAGTTTGAGCTG, reverse: CCTTGAGGCACACTTTGAAGTA; *Foxo1:* forward: *TCCAGTTCCTTCATTCTGCACT, reverse: GCGTGCCCTACTTCAAGGATAA; E-cadherín: forward: AGCTTTTCCGCGCTCCTG, reverse: CTTCCGAAAAGAAGGCTGTCC; Ghrelin: forward: GCCCAGCAGAGAAAGGAATCCA, reverse: GCGCCTCTTTGACCTCTTCC; Glucagon: forward: ATGAAGACCATTTACTTTGTGGCTG, reverse: CGGCCTTTCACCAGCCACGC; Lgr5: forward: ACCCGCCAGTCTCCTACATC, reverse: GCATCTAGGCGCAGGGATTG;* and *Ngn3: forward: TGGCCCATAGATGATGTTCG, reverse: AGAAGGCAGATCACCTTCGTG*. Using a Bio-Rad CFX96 real-time PCR system for measurements, relative gene expression was calculated using the ΔΔCt method with E-cadherin (as an epithelial marker of stomach cells) as the reference gene.

### Quantification

For quantification of tissue, at least at least 5 independent fields/samples for a minimum of 3 mice in each experimental group were quantified. Unless otherwise indicated, each dot in graphs of quantification of microscopic images represent an independent field. Number of specific mice for each experiment are indicated in figure legends. For primary cultures, stomach cells in 6–8 microscopic fields for at least 3 independent cultures were quantified.

### Statistics

A two-tailed student’s t-test was used to compare two groups. Analysis of variance followed by a Tukey post-hoc test Multiple comparisons was used to compare multiple groups. GraphPad Prism 6 software (La Jolla, CA) was used for calculations and P < 0.05 was considered significant. Graphs are depicted as average ± standard deviation with individual samples being indicated using individual points.

## Acknowledgements

We thank members of the Accili laboratory for insightful discussions and Tommy Kolar, Ana M. Flete-Castro, Xi Sun, and Lu Caisheng for technical support. This research was supported by a Berrie Fellow in Diabetes Research Award and 1K01DK121873 to W.M.M., DK103818 to U.P., and DK-57539 to D.A. These studies used the CCTI Flow Cytometry Core, supported by S10OD020056, and the resources of the Diabetes Research Center Flow Core, supported by P30DK063608.

## Author contributions

W.M.M. – study concept and design; acquisition of data; analysis and interpretation of data; drafting of the manuscript; obtained funding. S.S. – acquisition of data; analysis and interpretation of data. M.M. – acquisition of data; analysis and interpretation of data; critical revision of the manuscript for important intellectual content. M.K. – acquisition of data; analysis and interpretation of data. W.D. – analysis and interpretation of data. T.K. – analysis and interpretation of data. J.Y. – acquisition of data; analysis and interpretation of data. U.P. – analysis and interpretation of data; critical revision of the manuscript for important intellectual content. D.A. – study concept and design; analysis and interpretation of data; critical revision of the manuscript for important intellectual content; obtained funding.

